# High resolution kinematic approach for quantifying impaired mobility of dystrophic zebrafish larvae

**DOI:** 10.1101/2024.12.05.627004

**Authors:** Jeffrey J. Widrick, Matthias R. Lambert, Felipe de Souza Leite, Youngsook Lucy Jung, Junseok Park, James R. Conner, Eunjung Alice Lee, Alan H. Beggs, Louis M. Kunkel

## Abstract

Dystrophin-deficient zebrafish larvae are a small, genetically tractable vertebrate model of Duchenne muscular dystrophy well suited for early stage therapeutic development. However, current approaches for evaluating their impaired mobility, a physiologically relevant therapeutic target, are characterized by low resolution and high variability. To address this, we used high speed videography and deep learning-based markerless motion capture to develop linked-segment models of larval escape response (ER) swimming. Kinematic models provided repeatable, high precision estimates of larval ER performance. Effect sizes for ER peak instantaneous acceleration and speed, final displacement, and ER distance were 2 to 3.5 standard deviations less for dystrophin-deficient mutants vs. wild-types. Further analysis revealed that mutants swam slower because of a reduction in their tail stroke frequency with little change in tail stroke amplitude. Kinematic variables were highly predictive of the dystrophic phenotype with ≤ 3% of larvae misclassified by random forest and support vector machine models. Tail kinematics also performed as well as *in vitro* assessments of tail muscle contractility in classifying larvae as mutants or wild-type, suggesting that ER kinematics could serve as a non-lethal proxy for direct measurements of muscle function. In summary, ER kinematics can be used as precise, physiologically relevant, non-lethal biomarkers of the dystrophic phenotype. The open-source approach described here may have applications not only for studies of skeletal muscle disease but for other disciplines that use larval mobility as an experimental outcome.

## Introduction

Duchenne muscular dystrophy (DMD), a progressive pediatric neuromuscular disorder caused by out-of-frame mutations to the X-linked dystrophin gene (Koenig et al., 1987), is characterized by delays in motor development, excessive muscle weakness, impaired ambulation, and eventual cardio-respiratory failure (Guiraud et al., 2015). Developing therapies and cures for DMD has historically relied on small mammalian models, principally the *mdx* mouse. Dystrophin-deficient zebrafish larvae, such as *sapje* (Bassett et al., 2003) and *sapje-like* (Guyon et al., 2009), have emerged as alternatives or compliments to mammalian models due to their small size, rapid *ex utero* development, body transparency, permeability to small molecules, and genetic tractability. These features make zebrafish larvae ideal for probing the relationship between vertebrate genotype and disease phenotype and facilitate experimental approaches that would be difficult to conduct with other vertebrate DMD models, such as large-scale screens of potential therapeutics (Farr et al., 2020, Kawahara et al., 2011, Waugh et al., 2014).

High resolution approaches for quantifying phenotype are critical tools for the discovery and development of therapeutic compounds and treatments (Fuentes et al., 2018). However, there is growing concern in the broader biomedical research community that advances in genetics and molecular biology are outpacing the methods available for evaluating phenotype and that this imbalance may eventually impede scientific progress (Delude, 2015, Robinson, 2012). This is particularly important for zebrafish models of muscle disease where arguably their most physiologically relevant phenotype is mobility, yet current approaches for quantifying motor activity typically use low temporal and spatial resolution measurements of spontaneous motor activity (Widrick et al., 2023). Data collected in this manner are often characterized by large intra- and inter-larval variability while providing limited insight into mechanisms underlying impaired mobility.

The present project was designed to address these issues. Zebrafish larvae display a range of motor behaviors enabling them to hunt, capture prey, and avoid perceived threats (Budick and O’Malley, 2000, Burgess and Granato, 2007, Dunn et al., 2016, Müller and van Leeuwen, 2004). We chose to study their startle or escape response, a stereotypic behavior mediated by an all-or-none activation of the musculature of the tail and trunk that produces a brief burst of coordinated, high-intensity swimming activity (Franklin and Johnston, 1997, van Leeuwen et al., 2008, Wakeling and Johnston, 1998).

Because escape responses occur on a ms time scale, we utilized high speed videography to obtain high temporal resolution images of escape response movement. We then developed a markerless motion capture approach that extracted keypoint coordinates from the videos and enabled us to model each escaping larvae as a linked segment system. This approach revealed new insight into how the absence of dystrophin impacts vertebrate locomotion and identified statistically powerful, physiologically relevant, non-lethal biomarkers of the dystrophic phenotype.

## Methods

### Zebrafish

Zebrafish use was approved by the Institutional Animal Care and Use Committee at Boston Children’s Hospital (protocol #00001391). Adult heterozygous fish were mated. Fertilized eggs were collected and cultured in 100 cm petri dishes at 28.5°C with a 14 hr:10 hr light:dark cycle. All adult zebrafish and larvae were maintained in a fish facility overseen by the Aquatic Resources Program at Boston Children’s Hospital. Data on water quality, environmental conditions, diet, and other aspects of husbandry are available on protocols.io (https://dx.doi.org/10.17504/protocols.io.br4mm8u6).

### Birefringence assay

A non-lethal birefringence assay was used to classify larvae as affected or unaffected as previously described (Lambert et al., 2021, Smith et al., 2013). Briefly, lightly anesthetized larvae were aligned between two glass polarizing filters, viewed with a stereo-microscope, and classified as unaffected if the tail and trunk musculature appeared bright and well-organized or affected of the muscle displayed gaps, breaks, or loss of birefringence. Affected and unaffected larvae were then transferred into 48 well plates, one larvae per well, and returned to the incubator until further study.

### Experimental setup

Escape responses were evaluated in a rectangular arena formed by laser cutting a 20 mm x 30 mm opening in a 30 mm x 40 mm piece of acrylic. The arena was attached to the inside surface of a water-jacketed preparatory tissue dish (Radnoti, model 158401) with aquarium grade silicon. Water from a temperature-controlled bath circulated through the jacket and beneath the arena. The arena contained fish water to a depth of ≈ 3 mm at the center of the arena in order to minimize movements in the z-plane. A micro thermocouple confirmed that the fish water temperature was maintained at 25°C throughout data collection. An LED array and diffuser illuminated the arena from below.

### Escape responses

A single larva was transferred into the arena and given 5 min to temperature equilibrate. Escape responses were elicited by a 1 ms electric field pulse (Tabor et al., 2014) generated by a constant current muscle stimulator (Aurora Scientific, model 701) and delivered to platinum electrodes aligned along opposite walls of the arena. In preliminary studies, we established the current that consistently elicited an escape response and used this current for all subsequent trials.

Each larva was subjected to multiple trials until we had three acceptable escape responses (see quality control criteria below). A minimum of 60 s separated successive trials which is four times the inter-escape response period previously used to prevent habituation (Burgess and Granato, 2007).

### High speed videography

Videos (1280 x 864 pixels) were collected at 1000 frames/s using a monochrome high speed camera (Edgertronic SC2) positioned approximately 9 cm above the arena. A Nikon 50 mm f1.8D lens fitted with a 10X close-up lens produced a field of view that was roughly the same dimensions as the arena. Distance was calibrated each day of data collection and ranged from 44-45 pixels per mm.

A manually triggered, opto-isolated circuit was constructed to coordinate the escape response stimulus with the video recording. When triggered, the circuit opened the camera shutter but delayed the escape response stimulus by 10 ms producing 10 pre-stimulus frames for each video.

### Markerless pose estimation

The open-source machine learning toolkit DeepLabCut (version 2.2rc3) was used to create a deep neural network for estimation of larval pose (Mathis et al., 2018, Nath et al., 2019). To develop the neural network, we collected escape responses of 10 wild-type AB larvae (6 pdf). A kmeans algorithm selected 25 frames from each video that encompassed a diversity of poses. Each of the 250 frames were manually annotated with the following keypoints: TS, tip of the snout; S1, rostral swim bladder; S2, caudal swim bladder; T4 tip of trunk or tail; T2, point midway between S2 and T4; T3, point midway between T2 and T4; T1, point midway between S2 and T2. The keypoints defined six body segments: head, swim bladder, tail1, tail2, tail3, tail4. The keypoints were chosen so that we could quantify the length of the larvae’s body, track its approximate center of mass, and model the curvature of its body.

The neural network was initially trained using ResNet50 architecture for 7.5 × 10^5^ iterations using an 80% training to 20% testing split of the annotated dataset. Likelihood values were low when the tip of the tail was aligned very closely, to or even temporally occluded by, the head. To refine the network, we identified escape responses from six wild-type larvae where this occurred (3-8 frames per video). Keypoints with low likelihood scores (< 0.98) were either manually re-labeled or deleted (if occluded). These 28 newly annotated frames were merged with the original 250 annotated frames, the new data set was split into training (80%) and test (20%) sub-sets, and the neural network trained for 10^6^ iterations.

Network training was conducted on a Dell Precision 3640 desktop computer (Intel Core i7-10700K, 8 Core, 32 GB ram, Ubuntu version 20.04 LTS) equipped with a NVIDIA GeForce RTX 3080 graphical processing unit. With this configuration, the 10^6^ iterations used to train the neural network were completed in about 12 hours. Escape response videos were analyzed on the same hardware, requiring about 5 s per trial.

### Kinematic analysis overview

Custom scripts written in R (R Core Team, 2024) were used to calculate morphological and kinematic variables from the DNN output. Scripts used the following R packages: knitr (Xie, 2024), tidyverse (Wickham et al., 2019), data.table (Barrett et al., 2024), RcppRoll (Ushey, 2018), pracma (Borchers, 2023), patchwork (Pedersen, 2024), and kableExtra (Zhu, 2024).

### Angular kinematics

Tail segment angle data were smoothed using locally weighted smoothing (LOESS, span = 0.10) prior to analysis. Tail curvature was defined as the sum of the four individual trunk segment angles. Tail rotation direction was standardized between larvae by defining positive rotation as the direction of the first major bend of the tail (the C-start). Tail curvature angular velocity was calculated using the central finite difference equation (Robertson et al., 2014). The initiation of movement was determined as the post-stimulus frame where tail curvature exceeded the preceding frame by 2%.

### Linear kinematics

Linear kinematics were based on the movement of the larvae’s center of mass (COM). Larval COM is reported to fall at various locations between the rostral and caudal edges of the swim bladder (Budick and O’Malley, 2000, Müller et al., 2008, Müller and van Leeuwen, 2004, Stewart and McHenry, 2010). For the purposes of this study, we used keypoint S2 to represent the COM (Budick and O’Malley, 2000). Distance was defined as the length between S2 on consecutive frames. Displacement was defined as the vector between S2 position at movement time zero and the current S2 position. Distance and displacement were calculated from unfiltered data.

The instantaneous speed of S2 was determined as the first derivative of distance with respect to time using the central difference method. The acceleration of S2 was determined as the second derivative of displacement with respect to time. Instantaneous speed oscillates due to the undulatory nature of swimming. In order to increase consistency in determining the exit speed from each stage as well as the overall peak instantaneous speed, the raw speed response was smoothed using locally weighted smoothing (LOESS, span = 0.7). Linear kinematics were normalized to larval body length (BL). Body length was calculated as the sum of the 6 body segments, averaged across the pre-movement frames.

### Stage-specific kinematics

We calculated performance variables that took into account the entire escape response (overall distance, displacement, peak instantaneous speed, and peak acceleration). We also partitioned the escape response into three stages (Domenici and Hale, 2019) and determined stage specific kinematics. The first change in tail curvature sign delineated the transition from the stage 1 C-start to the stage 2 power stroke. The next change in sign indicated the end of the power stroke and the beginning of stage 3 burst swimming. Some larvae had a small counter tail bend that preceded the much larger C-start. These bends were ignored in analysis (Marques et al., 2018).

We used successive extremes in tail curvature to define a tail stroke. Thus, two tail strokes equals one tail beat cycle. Stage 1 and 2 consist of single tail strokes. Stage 3 consists of multiple strokes and we choose to carry the stroke nomenclature through stage 3 rather than convert strokes to tail beats. The number of stage 3 tail strokes varied between larvae. Therefore, for stage 3 we calculated variables that encompassed the entire stage (total stage duration, total stage distance, total stage displacement, and average stage speed) as well as variables that were based on the number of full strokes completed by the larvae, ignoring any partial strokes at the end of the stage. These variables were the average number of tail strokes, the average distance covered/tail stroke, the average tail stroke peak to peak amplitude, and the average tail stroke frequency.

### Muscle contractility

A subset of *sapje* strain larvae that had been subjected to ER analysis were subsequently used for assessment of muscle contractility as previously described (Widrick et al., 2016). The larvae was euthanized and one end of the tail was attached to the output tube of an isometric force transducer. Attachment was made at the gastrointestinal (GI) opening. This attachment point was easy to replicate as the 10-0 silk thread used for fastening the preparation was guided to this location by the notch formed at the intersection of the ventral and dorsal fin folds. This consistency in attachment enabled valid absolute force comparisons between preparations. The other end of the preparation was attached to an immobile titanium wire at a location that was approximately 2 mm distal to the GI opening.

Tail muscle preparations were studied in a bicarbonate buffer (equilibrated with 95% O_2_, 5% CO_2_) that was maintained at the same temperature as the escape response trials (25°C). A muscle twitch was induced using a single supra-maximal square wave pulse, 200 µs in duration, that was delivered to platinum electrodes flanking the preparation. The length of the preparation was adjusted to maximize twitch force. Contraction time, half-relaxation time, and the maximal rates of tension development and relaxation were calculated from the peak twitch force records (Widrick et al., 2016).

### Genotyping

Larvae were euthanized after completion of their escape trials. Genomic DNA was extracted from the heads and used as a PCR template. The *sapje* and *sapje-like* primer sets have been described by Spinazzola et al. (2020). Sanger sequencing was conducted at the Molecular Genetics Core Facility at Boston Children’s Hospital.

### Quality control

Raw videos were analyzed by the DNN without any pre-processing. Each escape response trial had to satisfy the following criteria in order to be included in further analysis. First, larvae had to remain stationary during the 10 ms pre-stimulus period. Second, the trial had to be accurately tracked by the neural network. Poor tracking was easily identified by a string of likelihood values < 0.98 coupled with erratic trunk curvature plots. Third, the video had to capture sufficient frames to enable us to analyze 60 ms of movement. Finally, only trials where there was agreement between fish phenotype (birefringence) and genotype (Sanger sequencing) were included in the final analysis, i.e. all affected fish genotyped as −/− and all unaffected fish as either +/+ or +/−.

### Statistical analysis

We evaluated 678 escape responses recorded from 49 *sapje* mutants and 73 wild-type siblings (29 +/+, 44 +/−) and 51 *sapje-like* mutants and 53 wild-type siblings (16 +/+, 37 +/−). These larvae were obtained from 7 and 4 clutches of *sapje* and *sapje-like* embryos respectively. Approximately 40% of the *sapje* strain larvae were subsequently used in the muscle physiology experiments (11 +/+, 20 +/−, 17 −/−). Unless otherwise noted, all results are based on these sample sizes.

Point estimates of effect sizes and variability were calculated using a bias-corrected-and-accelerated bootstrap approach. Standardized effect sizes were calculated using Hedges’s *g* with bias-corrected-and-accelerated intervals. Both methods used 5000 resamples and 99% confidence intervals.

To classify mutant (−/−) and wild-type (+/+ and +/−) groups, we employed a random forest model. All variables, regardless of stages, were used for modeling. For constructing trees, 2/3 of the data were used for training, and the remaining data were used for the out-of-bag calculation. To identify the important variables, the means of GINI importance were measured. To validate the random forest model, we employed a linear SVM model with various values of C (ranging from 0.00 to 5.00) to gain the most accurate model, allocating 70% of the data for training and 30% for testing. Our validation process included conducting a 10-fold cross-validation.

Statistical analysis, visualization, and graphics used R version 4.4.1 (R Core Team, 2024) and the following R packages: knitr (Xie, 2024), tidyverse (Wickham et al., 2019), cowplot (Wilke, 2024), patchwork (Pedersen, 2024), ggbeeswarm (Clarke et al., 2023), bootES (Kirby and Gerlanc, 2013), factoextra (Kassambara and Mundt, 2020), FactoMineR (Lê et al., 2008), ggh4x (van den Brand, 2024), pwr (Champely, 2020), ggtext (Wilke and Wiernik, 2022), randomForest (Breiman et al., 2024), ROCR (Sing et al., 2005), e1071 (Meyer et al., 2023), kernlab (Karatzoglou et al., 2004), ROCit (Khan and Brandenburger, 2024), gganimate (Pedersen and Robinson, 2024), circlize (Gu et al., 2014), and report (Makowski et al., 2023).

## Results

### Capturing escape responses

Initial studies were conducted on *sapje* zebrafish (Bassett et al., 2003) with follow-up studies conducted on the *sapje-like* strain (Guyon et al., 2009). Both mutants lack full-length dystrophin isoforms. Because mutants have a median lifespan of only 30 days (Bassett et al., 2003, Lambert et al., 2021, Spinazzola et al., 2020), larvae for study were obtained by breeding heterozygous adults (Figure 1-A). To facilitate studying dystrophin-positive and -negative larvae in a roughly 1:1 ratio (instead of an expected 3:1 ratio), we used a non-lethal birefringence assay to classify each larvae as having either an affected or unaffected muscle phenotype several days prior to their escape response trials.

**Figure 1.**
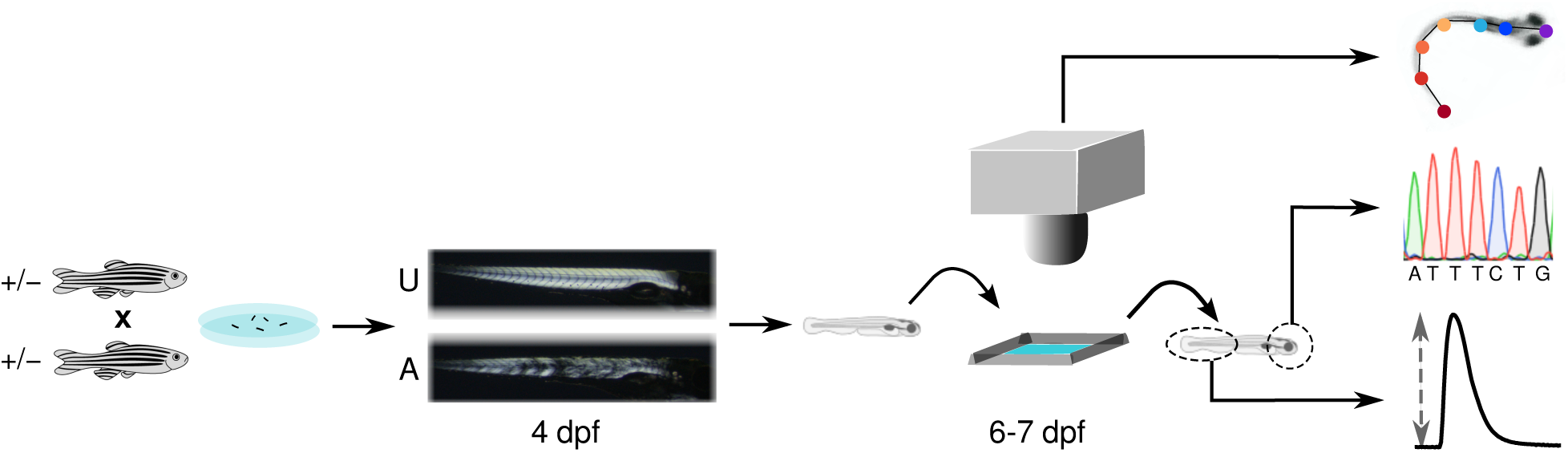
Overview of the experimental design. Larvae were obtained by mating heterozygous adults. At 4 day post-fertilization (dpf) a birefringence assay was used to classify larvae as having either an unaffected (U) or an affected (A) muscle phenotype. On 6 and 7 dpf, larvae were transferred one at a time into a small arena containing fish water where three escape responses, induced by a 1 ms electrical field pulse, were recorded at 1000 frames/s. Videos were used to evaluate linear and angular kinematics. Larvae were euthanized after their escape responses and DNA extracted from the head was used to confirm larval genotype. In a subset of post-escape response larvae, a portion of the tail was assayed for twitch contractile properties.

At 6 and 7 dpf, larvae were transferred one-at-a-time into an arena where escape responses were evoked with a brief electrical field pulse and recorded with high speed videography. Each recording consisted of a 10 ms pre-stimulus period, during which time the larvae was stationary, a variable duration latency period (≈7 ms) that separated the stimulus and the initiation of tail movement, and the subsequent escape swimming activity. We analyzed the initial 60 ms of escape response swimming. Longer periods of swimming were more difficult to collect as there was an increased likelihood that the larvae would exit the camera’s field of view before recording was complete. By 60 ms, most larvae had either reached their maximal speed or had attained a relatively constant speed.

Representative escape responses of a wild-type larvae and a *sapje* mutant are presented in Figure 2. For brevity, we only present selected time points in this Figure but even at this reduced resolution it is apparent that the dystrophic larvae swam considerably slower than expected. By the end of the 60 ms movement period the dystrophic larvae trailed its wild-type sibling by about 4 mm or 1 BL. Video 1 in the Supporting Information section is a video montage of these two escape responses in their entirety.

**Figure 2.**
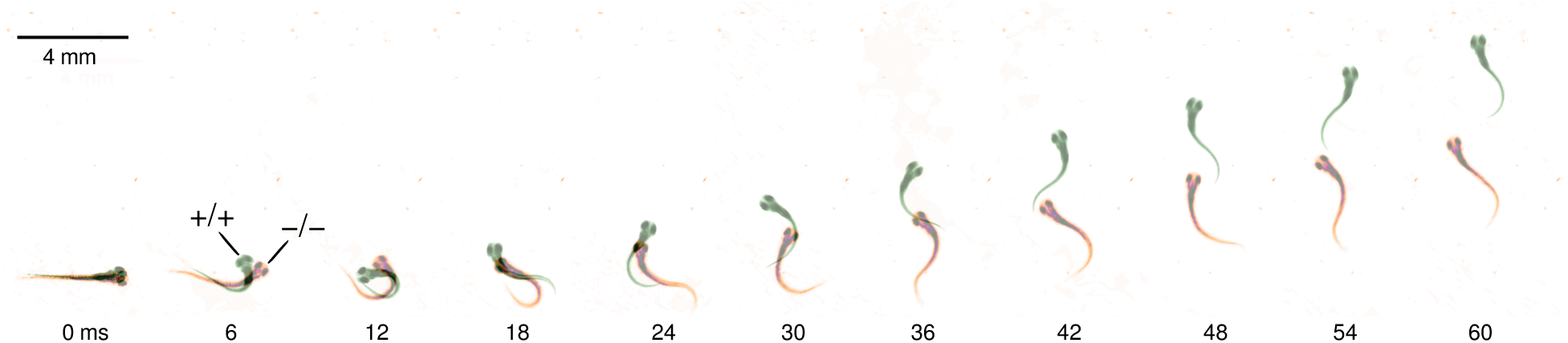
Representative escape responses. Shown are selected video frames (every 6th ms) from an escape response of a *sapje* mutant larva (−/−, pseudo-colored orange) and a wild-type sibling (+/+, pseudo-colored green). Numerals below the larvae indicate movement time in ms.

### A DNN accurately identifies keypoints on escaping larvae

In order to quantify escape response movements, we modeled each escaping larva as six linked body segments (Figure 3-A). We used the open-source pose estimation toolkit DeepLabCut (Mathis et al., 2018, Nath et al., 2019) to develop a deep-learning based neural network (DNN) for automated identification of the seven anatomical landmarks, or keypoints, that defined the body segments. The DNN was trained until it achieved a mean average Euclidean error (MAE) of 2.8 pixels which is equilibrant to 64 μm under our experimental conditions (Figure 3-B). This is similar to the test-retest accuracy of four human investigators tasked with manually identifying every third keypoint of a 60 ms escape response (Figures 3-C). In addition, one investigator manually annotated the complete escape response of a wild-type and a mutant larvae (over 800 total keypoints). The MAE between this manual annotation and the DNN estimates of the same two escape responses was 4.2 pixels (3-D). To place our MAE values in context, neural networks trained to identify keypoints on rodent models have reported MAE in the 2.0 to 4.7 pixel range (Ascona et al., 2024, Gonzalez et al., 2024, Sheppard et al., 2022).

**Figure 3.**
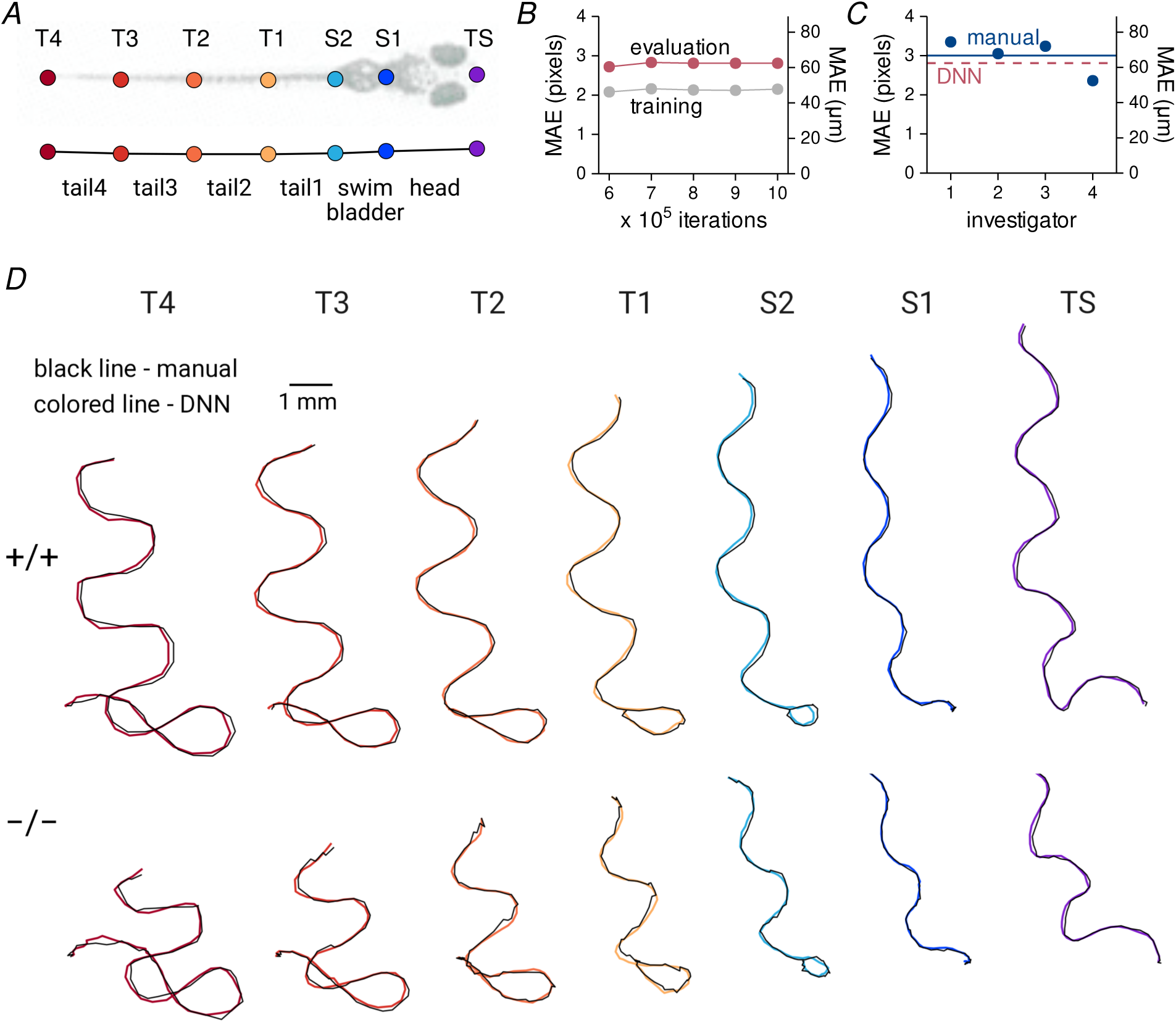
A deep neural network accurately estimates keypoints on escaping larvae. *A*, Keypoints and the corresponding body segments used for modeling larvae. *B*, The mean average Euclidean error (MAE) during training and evaluation of the deep learning-based neural network (DNN). *C*, Four investigators manually annotated portions of an escape response video on two occasions. The MAE between their initial trial and the repeat trial are shown as blue points and the average MAE is indicated by the solid blue line. For reference, the MAE of the DNN is indicated by the red dashed line. *D*, The most consistent annotator from Panel *C* manually annotated the two escape response videos shown in Figure 2. Keypoint escape response paths derived from the manual annotations and the DNN predictions are indicated by the black and colored lines, respectively.

### Modeling escape responses

The Cartesian coordinates output by the DNN were used to model each escaping larvae as a linked-segment system moving through 2-dimensional space. Figure 4-A shows the two larvae previously presented in Figure 2 now modeled as linked segments. A complete animated montage of these two models is available in the Supporting Information section (Video 2).

**Figure 4.**
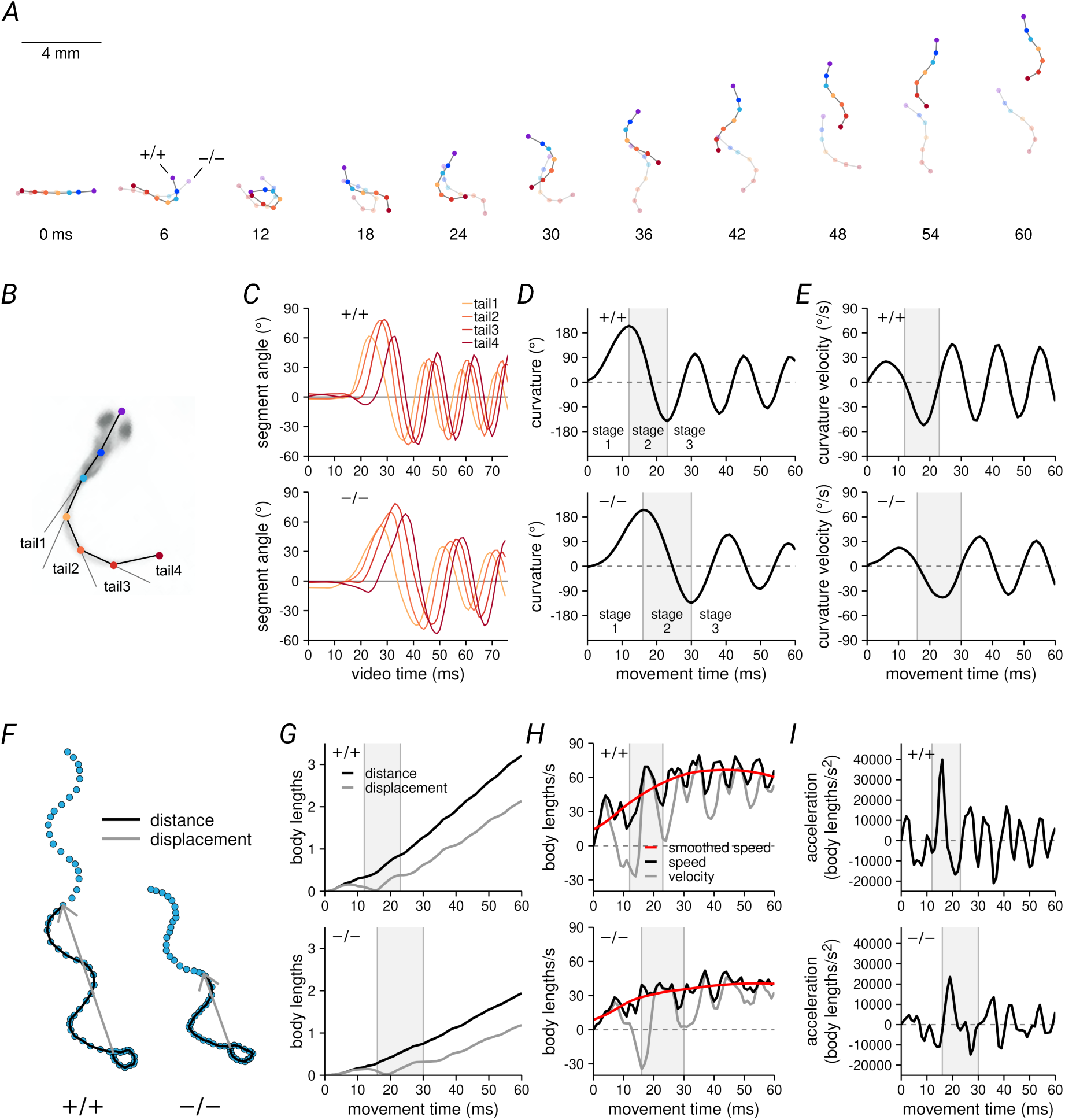
Quantification of escape response kinematics. *A*, Linked segment models of the two escape responses originally shown in Figure 2. All subsequent data in this figure derived from these two models. *B*, The coordinates of keypoints T1, T2, T3, and T4, were used to calculate the angle of each tail segment. *C*, Tail segment angles vs. movement time. *D*, Tail segment angle values were summed to yield tail curvature. Extremes in tail curvature was used to identify the transition from escape response stage 1 to 2 and from stage 2 to 3. *E*, Tail curvature angular velocity vs. movement time. *F*, Each point is the position of keypoint S2 (the putative center of mass) during the two escape responses. Distance and displacement at the 40th ms of movement are indicated by the solid black lines and gray vectors, respectively. *G*, Distance and displacement vs. movement time. *H*, Speed and velocity vs. movement time. Speed was smoothed for use in subsequent analyses. *I*, Acceleration was derived from velocity. Linear kinematics were normalized to larval body length.

From the keypoint coordinates, we calculated 31 kinematic variables that mathematically described the angular kinematics of the propulsive tail (Figure 4-B through -E) and the linear kinematics of the larva’s putative center of mass (Figure 4-F through -I). These variables could be categorized as those describing overall performance (60 ms distance and displacement and the peak instantaneous speed and acceleration attained during that time period) and those that quantified movement within three distinct biomechanical stages that make up the escape response (Domenici and Hale, 2019). These stages were defined as the initiating C-start, a high amplitude body bend that draws the head and tail together (stage 1), a rapid reversal in tail curvature that generates an initial power stroke (stage 2), and a concluding burst of tail strokes in rapid succession (stage 3).

### Repeatability of escape responses

We addressed the repeatability of kinematic measurements by focusing on the total distance covered in the response as this is a global marker of performance that encapsulates all other kinematic variables. Each larvae completed three escape trials. In general, responses were consistent within a genotype, across days, and for both strains (Figure 5). To quantify this apparent consistency, we calculated trial to trial repeatability coefficients (Bland and Altman, 1996). For repeated distance measurements made on the same larvae, trial-to-trial differences would be expected to exceed 0.37 BL in only 5% of cases.

**Figure 5.**
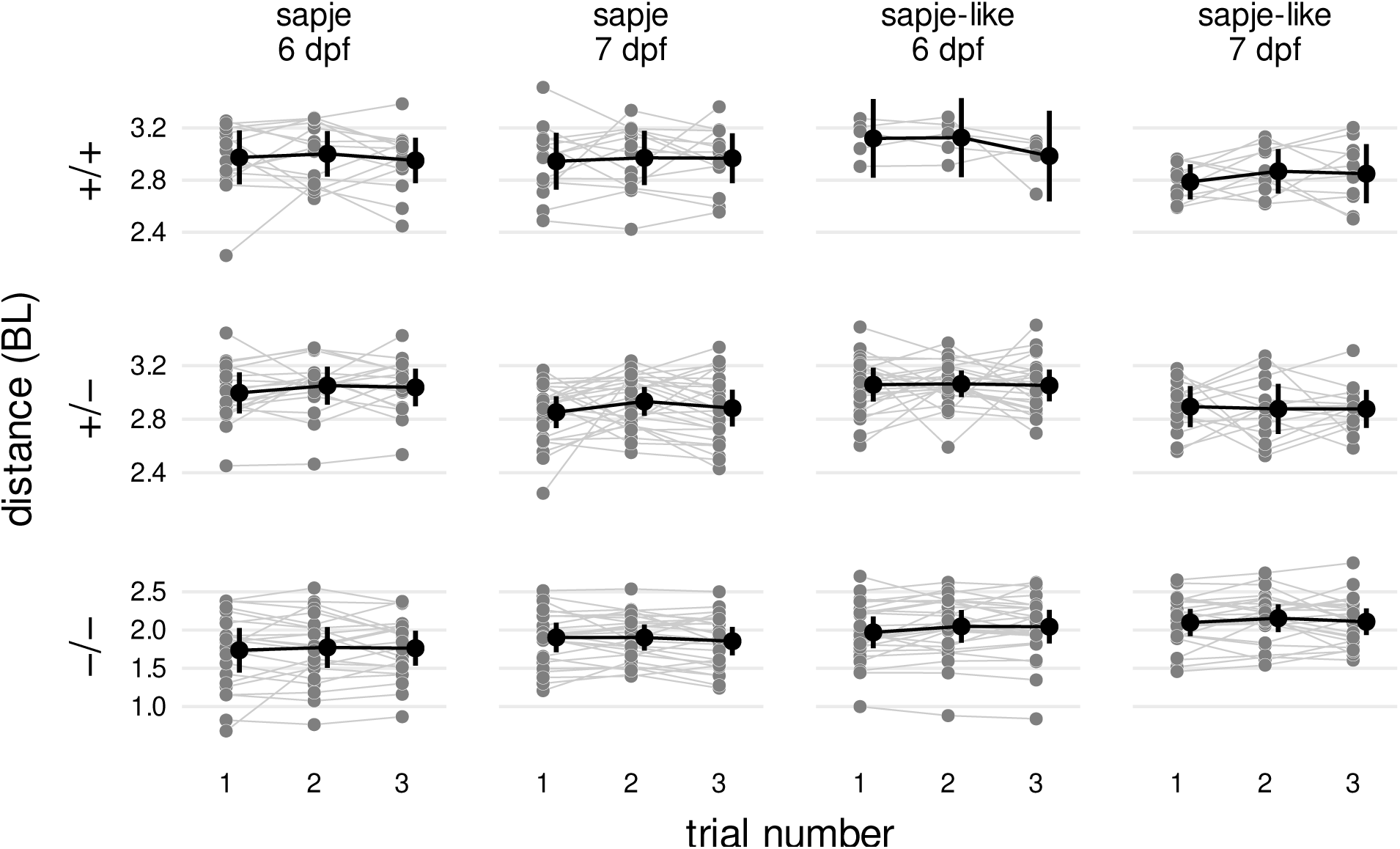
Escape response distance measurements are repeatable. Each gray symbol represents the distance covered during a single escape response trial. Trials from the same larvae are connected with lines. Black points are the trial mean and 99% confidence interval. Abbreviations: BL, body length. Data from 366 videos of *sapje* larvae (122 larvae, 3 trials/larvae) and 312 videos of *sapje-like* larvae (104 larvae, 3 trials/larvae).

Based on the repeatability of this global indicator of overall escape performance, the three trials were collapsed into an average value per larvae for all variables. All data points and analyses going forward are based on a data set where each variable is represented by a single mean value per larvae.

### Impaired escape response performance of dystrophic larvae

We used escape response distance, final displacement, peak instantaneous speed, and peak acceleration as markers of overall escape response performance. The dot plots in the top row of Figure 6-A and -B show performance values for each individual *sapje* and *sapje-like* larvae along with the group means and 99% confidence intervals. The confidence intervals can be interpreted as the measurement error and are indicative of the high precision of the kinematic measurements.

**Figure 6.**
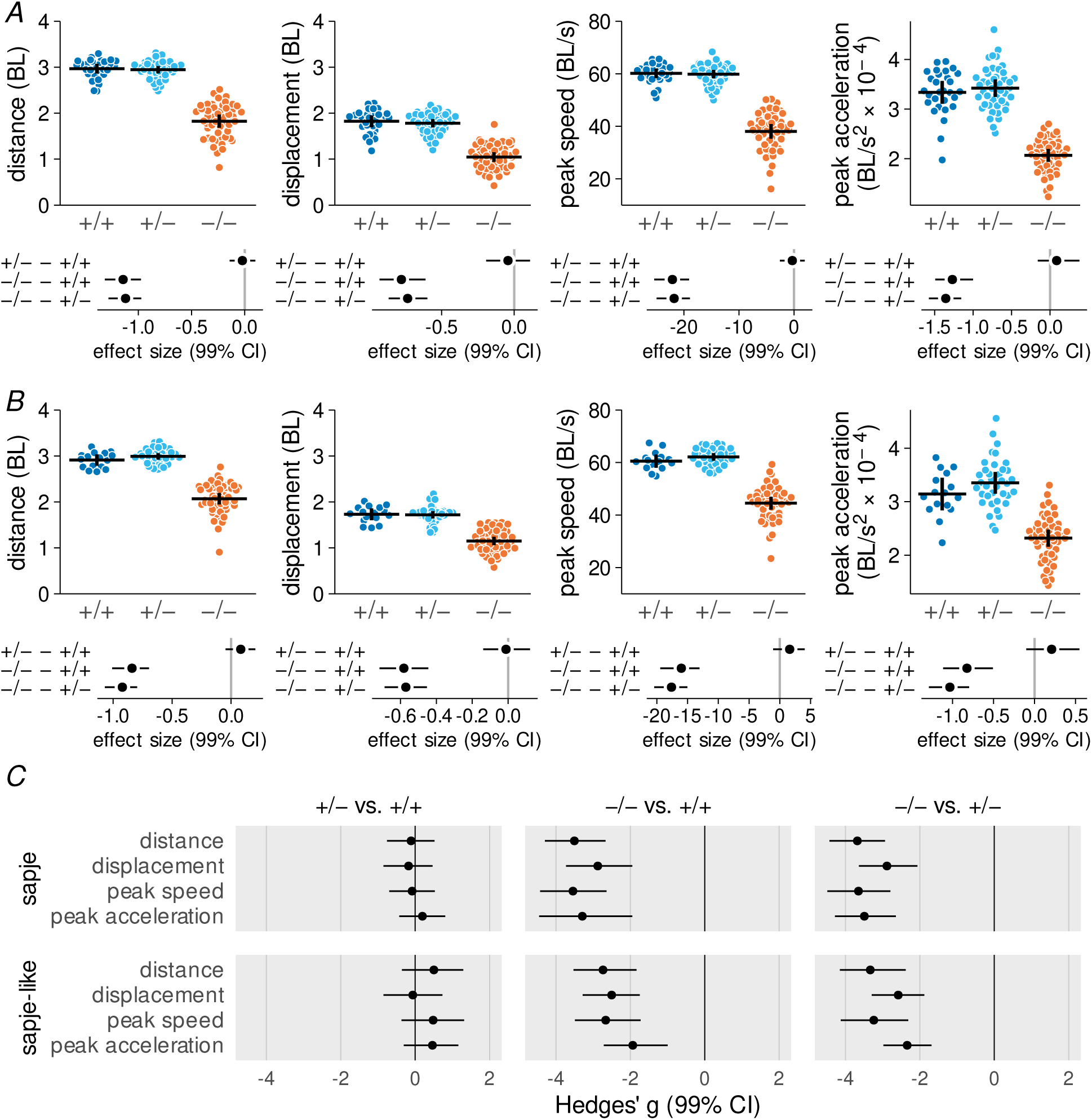
Escape response performance is impaired in dystrophic larvae. *A*, *sapje* larvae. *B*, *sapje-like* larvae. In Panels *A* and *B* each data point in the upper row of plots represents the mean value for a single larva (mean of three separate trials) with the horizontal bars indicating the genotype group mean and 99% confidence intervals. Below the dot plots are the bootstrapped effect sizes with 99% confidence intervals for each pairwise comparison (in the same units as the dependent variable). *C*, Bootstrapped Hedges’ *g* with 99% confidence intervals for each genotype contrast. Units are standard deviations. Abbreviations: BL, body length. Data from 122 *sapje* strain larvae (29 +/+, 44 +/−, 49 −/−) and 104 *sapje-like* strain larvae (16 +/+, 37 +/−, 51 −/−).

Several kinematic variables did not appear to be normally distributed. To address this, we used a bootstrap procedure to re-sample the data set and calculate bootstrapped means and 99% confidence intervals for all between group comparisons (Kirby and Gerlanc, 2013). These bootstrapped values are presented immediately below their respective dot plot in Figure 6-A and -B. Based on the bootstrapped results, there was little evidence that escape response performance differed between wild-type homozygous and heterozygous larvae. In contrast, *sapje* and *sapje-like* mutants consistently under-performed their dystrophin-positive siblings across all four variables presented here. Taking escape response distance as an example, our best estimate is that *sapje* mutants covered 1.1 BL less distance after 60 ms of escape response swimming compared to the dystrophin-positive larvae, with plausible values ranging from a deficit of 0.98 to a deficit of 1.3 BL (plausible values are defined by the 99% confidence intervals). Because a between group difference of zero is not a plausible value, the differences between mutants and the wild-type groups are statistically significant (at p < 0.01 or less).

Mutants also displaced themselves less than expected during an escape response but effect sizes for displacement were about one-third less than noted for distance. Comparisons between distance and displacement are possible because both are expressed in the same units. In order to facilitate comparisons of effect sizes across variables of different measurement units, we calculated bootstrapped Hedges’s *g* (Hedges, 1981), a standardized effect size that is expressed in units of standard deviations (SD) (Figure 6-C). Bootstrapped effect sizes of 3.5 SD distinguished distance, peak instantaneous speed, and peak acceleration of *sapje* mutants from their wild-type siblings, with the effect size for displacement slightly less at 2.8 SD. For *sapje-like*, effect sizes were somewhat less than observed in *sapje* larvae but were still substantial, ranging from 1.9 to 3.3 SD’s.

### Biomechanical mechanisms underlying impaired mobility

To identify reasons why escape response swimming performance was impaired in dystrophic larvae we evaluated 27 kinematic properties that quantified center of mass movement and tail mechanics. Figure 7 presents Hedges’ *g* for these kinematic variables classified by escape response stage for both strains of larvae. The left most column of plots indicates that there is little difference in the escape response kinematics of wild-type homozygous and heterozygous larvae. The second and third columns reveal that *sapje* and *sapje-like* mutants show a very similar escape response kinematic profile and that in each case, these profiles differ markedly from their respective wild-type siblings.

**Figure 7.**
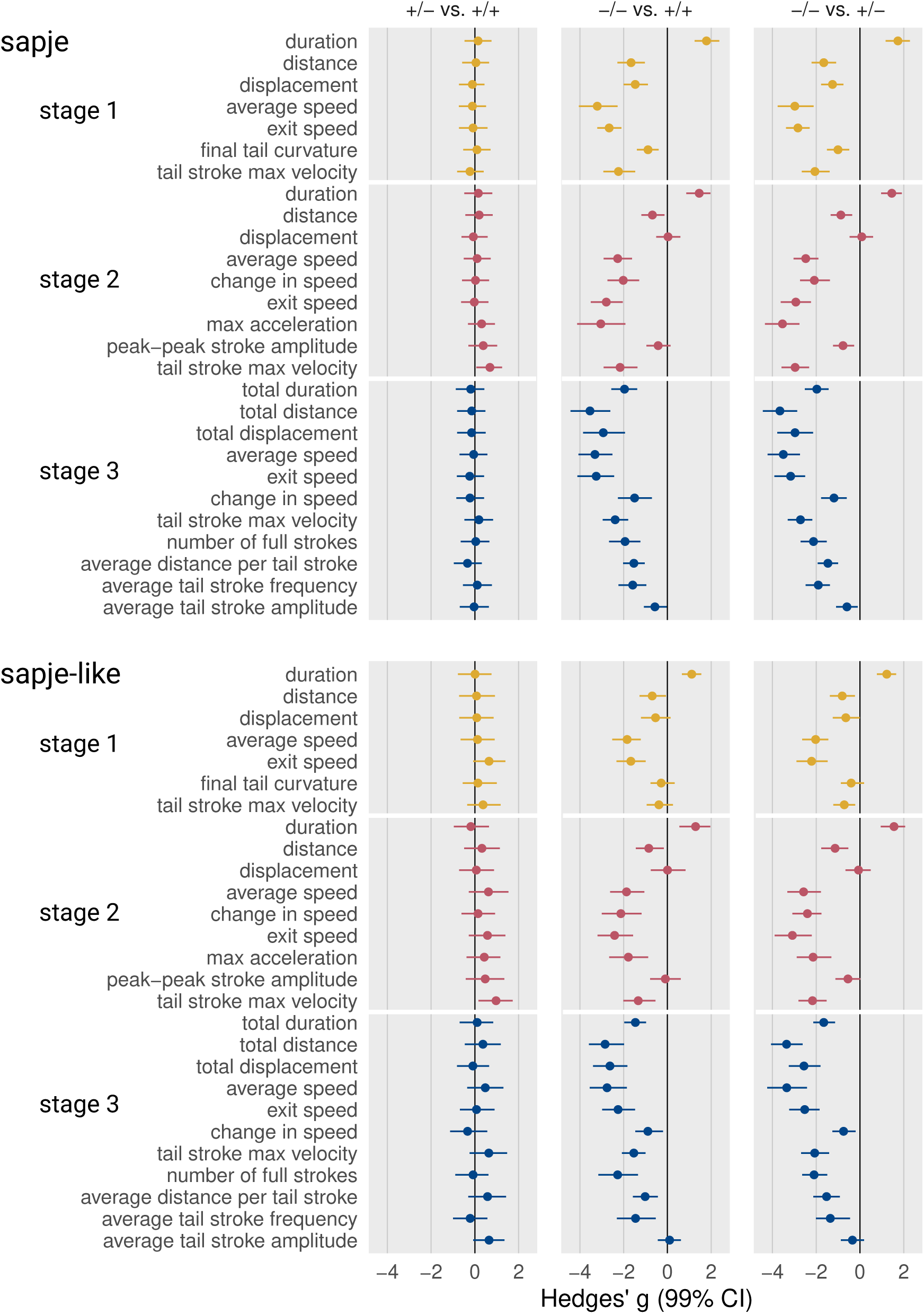
Kinematic profiles reveal mechanisms underlying impaired swimming of dystrophic larvae. Points are bootstrapped Hedges’ *g* with 99% confidence intervals. Units are in standard deviations. Plot interpretation and sample sizes same as Figure 6-C. Abbreviations: max, maximum.

Larval swimming speed is a function of the amplitude and frequency of tail strokes and Figure 7 provides insight into how these parameters were impacted by the absence of dystrophin. Peak to peak tail amplitude appeared to be only mildly affected, if affected at all, in dystrophin mutants. For instance, there were relatively small differences between mutant and wild-type larvae in the final tail curvature of stage 1, the change in tail curvature during the power stroke of stage 2, and the average amplitude of tail strokes in stage 3. In contrast, mutants showed a substantial slowing in the maximum angular velocity of tail strokes across all three stages and a slowing of stage 3 average tail stroke frequency.

A slower oscillating tail, even when coupled with a roughly unaffected tail stoke amplitude, will produce less propulsive power to move the larvae forward. Thus, the slower stage 2 power stroke of mutants generated less acceleration (stage 2 max acceleration) which slowed the larvae’s speed as it began stage 3 burst swimming (stage 2 exit speed). Then in stage 3, the distance that each tail stroke propelled the dystrophic larvae (stage 3 average distance/tail stroke), the average speed of the larvae (stage 3 average speed) and its speed at the 60 ms time point (stage 3 exit speed) were all reduced, contributing to a reduction in the distance the larvae covered in stage 3 (stage 3 distance).

In addition to impacting propulsion, slower tail angular velocities and stroke frequencies affected several other aspects of the escape response that exacerbated the decline in performance. Slower tail strokes prolonged stages 1 and 2 which in turn left less time for stage 3 swimming. Because stage 3 swimming time was reduced, dystrophic larvae completed fewer stage 3 tail strokes (stage 3 average number of tail strokes) than wild-type larvae. These changes were critical as the majority of the distance covered in 60 ms of escape response swimming (see Figure 4-G) normally takes place in stage 3.

### Kinematics are predictive of the dystrophic genotype

We next asked whether kinematic variables could be used to predict larval genotype. A random forest model, using all 31 kinematic variables, was developed to differentiate between dystrophic (−/−) and wild-type larvae (+/+ and +/−) of both the *sapje* and the *sapje-like* strains. The model discriminated between mutants and wild-types with high accuracy, attaining AUROC (area under the receiver operating characteristic curve) values of 0.994 to 0.999 and error rates of ≤ 2.5% (Figure 8-A). We confirmed these results using a second predictive model, a support vector machine (SVM). The SVM analysis was equally accurate, differentiating mutant from wild-type larvae with an accuracy of 97% for the *sapje* strain and 1.00% for the *sapje-like* strain (Figure 8-B).

**Figure 8.**
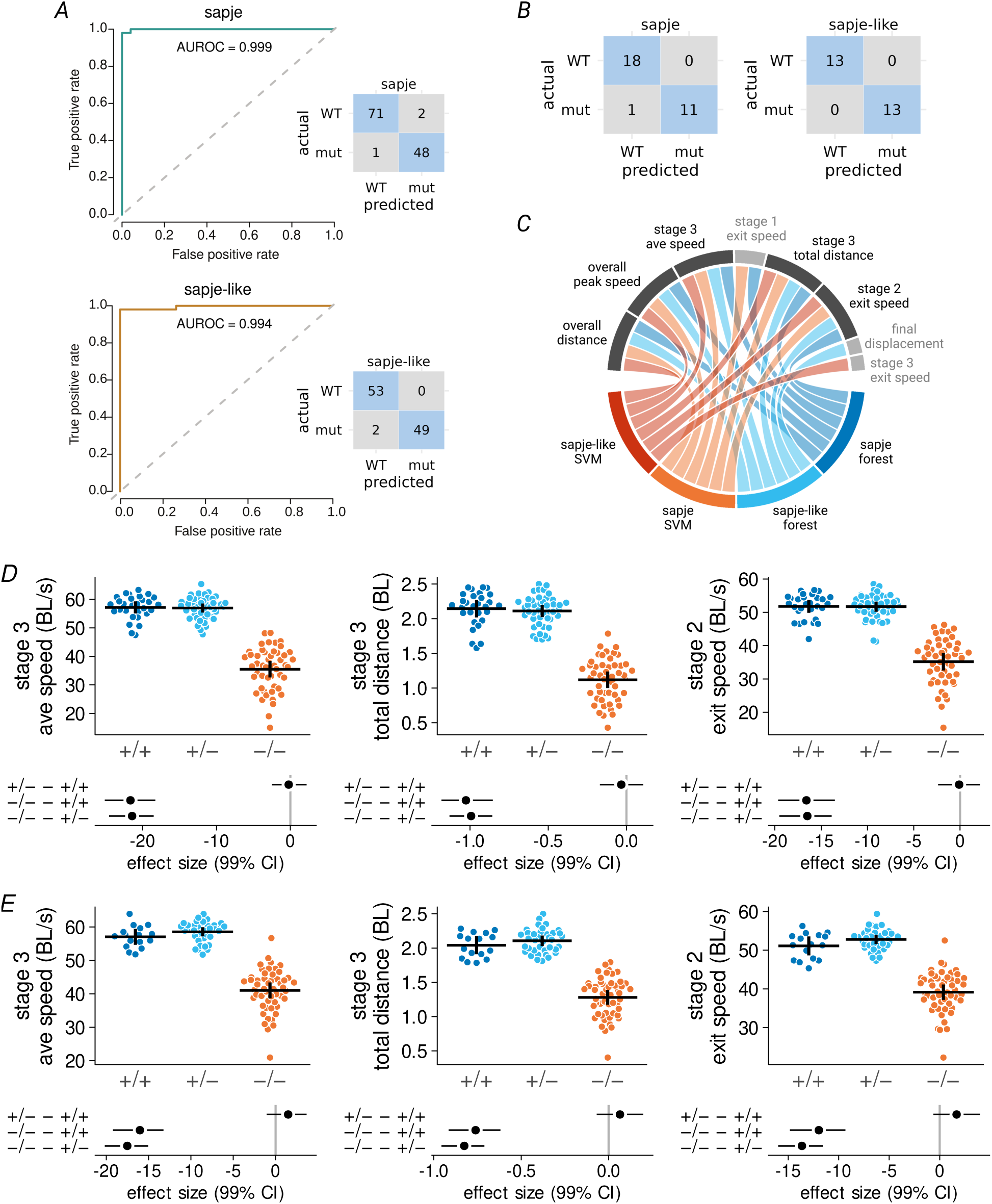
Swimming kinematics are highly predictive of larval genotype. *A*, Random forest model area under receiver operating characteristic curve and confusion matrix results for *sapje* and *sapje-like* larvae. *B*, Support vector machine (SVM) model confusion matrices for *sapje* and *sapje-like* larvae. *C*, Chord plot of the 6 best predictors from the forest and SVM classification models. Five kinematic variables were identified that were common to all four of the strain by predictor model combinations. Descriptions of two of these variables (overall distance and overall peak speed) have been presented in Figure 6. The remaining variables are presented in detail here for *D*, *sapje* larvae, and *E sapje-like* larvae. Interpretation and samples sizes of Panel *D* and *E* same as in Figure 6-A and -B, respectively. Abbreviations: mut, mutant; AUROC, area under receiver operating characteristic curve.

In order to identify which of the 31 kinematic variables were most predictive, we created a chord plot based on the top six predictors from each of the four strain by predictive model combinations (Figure 8-C). This plot revealed that five kinematic variables were common to each combination: overall distance and peak instantaneous speed (previously discussed in Figure 6), the average speed during stage 3 burst swimming and the distance covered in that stage, and the speed with which larvae exited stage 2 following the power stroke (Figure 8-D and -E). These five variables capture key kinematic features of the escape response that distinguish dystrophin-deficient mutants from wild-type larvae.

### Kinematics predict genotype as well as measurements of muscle contractility

Dystrophic zebrafish larvae are characterized by weak tail muscles (Li et al., 2014, Widrick et al., 2016) and we asked if this weakness was consistent with their impaired escape response swimming. We confirmed tail muscle weakness in sub-groups of mutant and wildtype *sapje* larvae after we had collected videos of their escape response performance. *In vitro* tail muscle preparations indicated that preparations from mutant *sapje* larvae generated only 42% of the twitch force observed for preparations from wild-type larvae (Figure 9-A and -B). Twitch contraction time and half-relaxation time were similar across genotypes but the rates of twitch tension development and relaxation were reduced for the *sapje* mutants (Figure Figure 9-C to -F).

**Figure 9.**
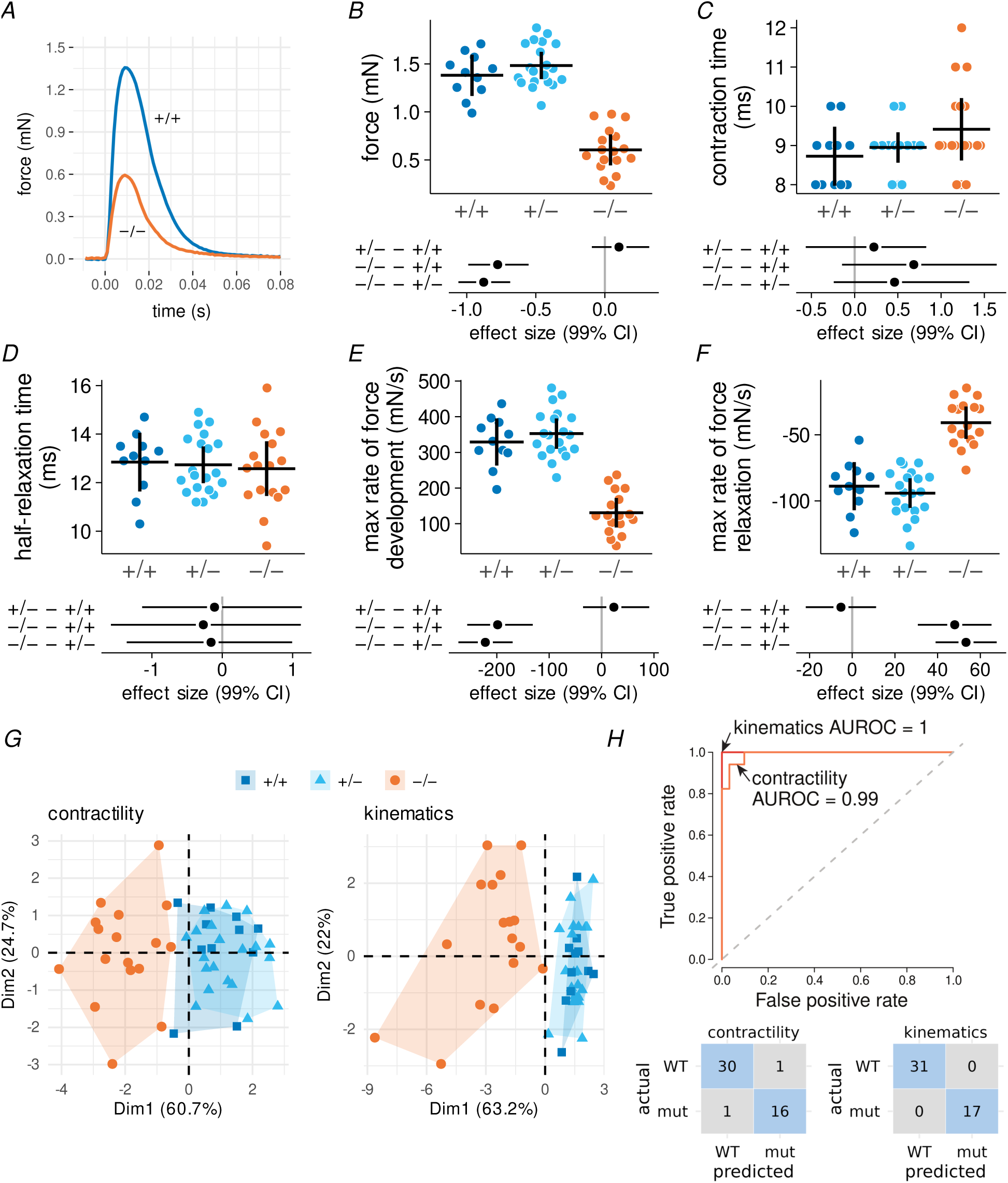
Tail angular kinematics and tail muscle contractility predict genotype with similar accuracy. *A*, Representative twitch force records from tail muscle preparations of a wild-type and *sapje* mutant. Twitches were used to determine *B*, peak force, *C*, contraction time, *D*, half-relaxation time, *E*, maximum rate of force development, and *F*, maximum rate of force relaxation. Sample sizes for contractility measurements: 11 +/+, 20 +/−, 17 −/− *sapje* larvae. *G*, Biplot of principle components 1 (Dim1) and 2 (Dim2) for these five contractility variables and nine variables from Figure 7 describing tail kinematics (see text for details). *H*, Random forest classification models for the prediction of genotype using contractile or kinematic variables. Abbreviations: mut, mutant; AUROC, area under receiver operating characteristic curve.

We tested how well these five measures of tail muscle contractile function differentiated between genotypes. We did the same for nine kinematic variables describing the motion of the propulsive tail (from stage 1: duration (which is the same as tail stroke duration), final tail curvature, and tail stroke maximum angular velocity; from stage 2: tail stroke duration, tail stroke amplitude, and tail stroke maximal angular velocity; from stage 3: average tail stroke frequency, average tail stroke amplitude, and tail stroke maximal angular velocity). Principal component analyses indicated that the first two dimensions accounted for 80% of the total variability between genotypes regardless of whether the analysis was conducted with contractile or kinematic variables and that both approaches appeared equally capable of segregating dystrophic from wild-type larvae (Figure 9-G). A random forest model using muscle contractility as a predictor misclassified the genotype of 2 out of 48 larvae (Figure 9-H). The same model using kinematic variables as predictors correctly classified all larvae.

## Discussion

Evoking escape responses by gentile tactile stimulation and visually judging if swimming is impaired has been used for many years as an easily administered, but subjective, way to identify mobility mutants in large-scale mutagenesis screens (Granato et al., 1996, Gupta et al., 2013, Smith et al., 2013, Telfer et al., 2012). While these touch-evoked escape responses are a valuable tool for identifying skeletal muscle mutants, the approach lacks the temporal and spatial resolution necessary for quantifying locomotor dysfunction. The present study updates and extends the classic touch-evoked approach by using high speed data acquisition to improve temporal resolution, machine learning to facilitate modeling escaping larvae as linked-body segments, and kinematic evaluation to quantify swimming performance and the biomechanical mechanisms responsible for locomotion.

We observed large effect size deficits in escape response swimming of dystrophin mutants. Following the first 60 ms of escape response movement, mutants trailed their wild-type siblings by an average of over 1 body length. By examining the mechanics of the propulsive tail, we found that the absence of dystrophin had little or a very modest effects on peak-to-peak tail stroke amplitude during an escape. In contrast, there was a substantial slowing in the frequency and angular velocity of tail strokes. Slower tail strokes would be expected to have a direct effect on the generation of power for propulsion, consistent with the reduced acceleration produced by the stage 2 power stroke and the subsequent slower swimming speed of mutant larvae. A slowing of tail strokes also had a cascading effect on temporal aspects of our 60 ms escape responses, such as reducing the time available for critical stage 3 burst swimming.

While a slowing in the oscillating motion of the tail seems to be the primary mechanisms underlying the impaired mobility of dystrophin mutants, we found no evidence that the intrinsic rate of muscle contraction was altered in isolated tail muscle preparations from mutant *sapje* larvae. For example, we detected no difference in twitch contraction time between mutant and wild-type tail muscle preparations. In the absence of evidence for a slowing in intrinsic shortening, we propose that the slowing of the tail stroke rate is a secondary consequence of the muscle weakness that characterizes dystrophin mutants.

We identified five kinematic variables that were highly predictive of the dystrophic genotype. These predictors appear quite robust as all five were identified by two independent prediction models applied to two different strains of dystrophic larvae. These variables could be valuable biomarkers of the dystrophic genotype in future studies. All five were linear kinematic measurements: the overall distance the larvae swam in 60 ms, its peak instantaneous speed during that time, the distance covered and average speed during stage 3 swimming, and the stage 2 exit speed. It is important to note that because zebrafish are undulatory swimmers, high sampling rates are required in order to quantify the distance they cover during a swimming bout. Measurements made from where a larvae starts moving to where it stops defines their displacement and displacement measurements may not be as robust as the candidate biomarkers identified here.

Because of the precise nature of these biomarkers, they are also statistically powerful. To illustrate, we calculated theoretical statistical power curves based on the present data (Figure 10). These curves show that if a treatment improves escape response distance of a group of dystrophin mutant larvae by a modest 20%, this difference would be detectable (80% power at p = 0.05) from an untreated group of mutants with sample sizes of about 15 larvae/group. Because our kinematic approach is non-lethal, studying the same larvae before and after treatment may be possible. In this paired experimental design, the required sample size falls to less than 10 larvae. These sample sizes are considerably less than the hundreds of larvae per condition that are sometimes used when assessing spontaneous swimming activity (Lambert et al., 2021).

**Figure 10.**
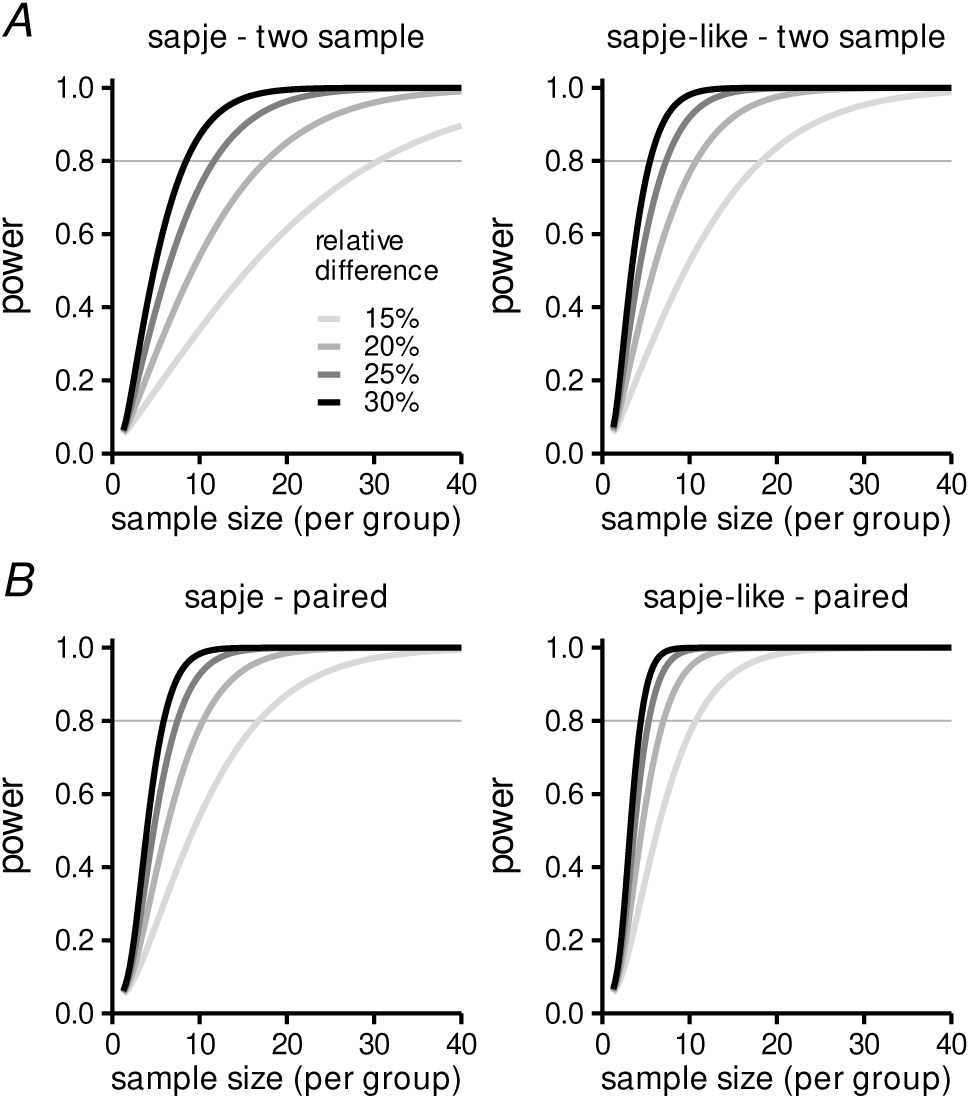
Swimming kinematics have high estimated statistical power. Estimates are based on detecting relative differences in escaped response distance ranging from 15% to 30% using a two sided t-test at p = 0.05. *A*, Power curves for a two-sample design. *B*, Power curves for a paired design. Power curves were calculated using the R package ‘pwr’ (Champely (2020)) and were based on the mean and SD of the *sapje* and *sapje-like* mutant larvae in this project and the assumption that the SD’s do not change as a result of treatment.

Tail kinematics and muscle contractility accounted for almost identical amounts of variability in our data set and both approaches were excellent predictors of larval genotype. These findings support the use of kinematic measurements as a proxy for direct *in vitro* measures of tail muscle contractility. Compared to *in vitro* measurements, kinematic analysis presents a number of advantages: it is non-contact, non-lethal, and technically less demanding. Its non-lethal nature is particularly noteworthy as it opens up new possibilities in experimental design, such as serial measurements conducted on the same larvae.

The kinematic approach outlined here has certain limitations. First, as currently configured, it is not a high throughput approach. This is due to biological factors such as the recovery time between trials to prevent fatigue and habituation, and technical factors which stem mainly from the small size of larvae and their tendency to spontaneously swim out of the cameras field of view. While the present approach is not amenable to high throughput screening, it can serve as a way to more thoroughly evaluate candidate compounds before they are advanced to more resource-intensive mammalian models.

A second limitation concerns fiber type recruitment. The musculature of the larval tail consists of a thin, superficial layer of slow contracting red fibers surrounding a vastly larger volume of fast, white fibers (Devoto et al., 1996, van Raamsdonk et al., 1982). Both slow and fast fibers are recruited during a larval escape response (Buss and Drapeau, 2000) but because of the sheer volume and faster contraction speed of the white fibers, the fast motor unit pool generates the propulsive power for movement (van Leeuwen et al., 2008). This is an advantage when using escape responses to evaluate mutations that impact all fiber types or fast fibers exclusively but presents a limitation when studying disorders that impact the small pool of slow, oxidative fibers. Preliminary data from our laboratory indicates that escape response kinematics may fail to identify larvae with mutations impacting mitochondrial function (Barraza-Flores et al., 2024) or that target proteins expressed exclusively in slow muscle fibers (Lambert and Gussoni, 2023). Investigators may need to consider other approaches for evaluating mobility when studying fish with these types of mutations.

In summary, we have developed high temporal and spatial resolution kinematic models of escaping zebrafish larvae. From the models we identified repeatable, precise, physiologically relevant kinematic variables that reveal large effect size differences in the swimming performance of dystrophin-deficient and wild-type larvae. The approach described here opens new possibilities for future studies using zebrafish larvae to investigate DMD, other muscle diseases, and movement disorders in general. More broadly, this approach may be useful in any discipline that uses high-intensity motor activity of zebrafish larvae as an experimental outcome.

## Acknowledgments

The authors are grateful to Pamela Barraza-Flores for sharing her insight into larvae with mitochondrial defects and to Sheldon Oliveira for his assistance with zebrafish husbandry. Genotyping was conducted by the Boston Children’s Hospital IDDRC Molecular Genetics Core Facility which is funded by P50HD105351 from the National Institutes of Health of USA.

## Author contributions

Conceptualization: JJW, MRL, FSL, AHB, LMK

Methodology: JJW, JRC

Investigation: JJW, MRL, FSL, JRC

Data analysis: JJW, YLJ, JP, EAL

Software development: JJW Visualization: JJW

Supervision: JJW, EAL, AHB, LMK

Writing—original draft: JJW Writing—review & editing: all authors

## Supporting Information

Video 1. Video montage of the two escape responses shown in Figure 2.

Video 2. Animated montage of the two escape responses modeled in Figure 4-A.

## References

Ascona MC, Tieu EK, Gonzalez-Vega E, Liebl DJ & Brambilla R (2024). A deep learning-based approach for unbiased kinematic analysis in CNS injury. Exp Neurol 381, 114944.

Barraza-Flores P, Moghadaszadeh B, Lee W, Isaac B, Sun L, Troiano EC, Rockowitz S, Sliz P & Beggs AH (2024). Zebrafish and cellular models of SELENON-Related Myopathy exhibit novel embryonic and metabolic phenotypes.

Barrett T, Dowle M, Srinivasan A, Gorecki J, Chirico M & Hocking T (2024). data.table: Extension of ‘data.frame‘ https://CRAN.R-project.org/package=data.table.

Bassett DI, Bryson-Richardson RJ, Daggett DF, Gautier P, Keenan DG & Currie PD (2003). Dystrophin is required for the formation of stable muscle attachments in the zebrafish embryo. Development 130, 5851–5860.

Bland JM & Altman DG (1996). Measurement error. BMJ 313, 744.

Borchers HW (2023). pracma: Practical numerical math functions https://CRAN.R-project.org/package=pracma.

Breiman L, Cutler A, Liaw A & Wiener M (2024). randomForest: Breiman and Cutlers random forests for classification and regression https://cran.r-project.org/package=randomForest.

Budick SA & O’Malley DM (2000). Locomotor repertoire of the larval zebrafish: swimming, turning and prey capture. J. Exp. Biol. 203, 2565–2579.

Burgess HA & Granato M (2007). Sensorimotor gating in larval zebrafish. J. Neurosci. 27, 4984–4994.

Buss RR & Drapeau P (2000). Physiological properties of zebrafish embryonic red and white muscle fibers during early development. J. Neurophysiol. 84, 1545–1557.

Champely S (2020). pwr: Basic functions for power analysis https://CRAN.R-project.org/package=pwr.

Clarke E, Sherrill-Mix S & Dawson C (2023). ggbeeswarm: Categorical scatter (violin point) plots https://CRAN.R-project.org/package=ggbeeswarm.

Delude CM (2015). Deep phenotyping: The details of disease. Nature 527, S14–15.

Devoto SH, Melançon E, Eisen JS & Westerfield M (1996). Identification of separate slow and fast muscle precursor cells in vivo, prior to somite formation. Development 122, 3371–3380.

Domenici P & Hale ME (2019). Escape responses of fish: a review of the diversity in motor control, kinematics and behaviour. J. Exp. Biol. 222, 1–15.

Dunn TW, Mu Y, Narayan S, Randlett O, Naumann EA, Yang CT, Schier AF, Freeman J, Engert F & Ahrens MB (2016). Brain-wide mapping of neural activity controlling zebrafish exploratory locomotion. Elife 5, e12741.

Farr GH, Morris M, Gomez A, Pham T, Kilroy E, Parker EU, Said S, Henry C & Maves L (2020). A novel chemical-combination screen in zebrafish identifies epigenetic small molecule candidates for the treatment of Duchenne muscular dystrophy. Skelet Muscle 10, 29.

Franklin CE & Johnston IA (1997). Muscle power output during escape responses in an Antarctic fish. J Exp Biol 200, 703–712.

Fuentes R, Letelier J, Tajer B, Valdivia LE & Mullins MC (2018). Fishing forward and reverse: Advances in zebrafish phenomics. Mech Dev 154, 296–308.

Gonzalez M, Gradwell MA, Thackray JK, Patel KR, Temkar KK & Abraira VE (2024). Using DeepLabCut-Live to probe state dependent neural circuits of behavior with closed-loop optogenetic stimulation.

Granato M, van Eeden FJ, Schach U, Trowe T, Brand M, Furutani-Seiki M, Haffter P, Hammerschmidt M, Heisenberg CP, Jiang YJ, Kane DA, Kelsh RN, Mullins MC, Odenthal J & Nüsslein-Volhard C (1996). Genes controlling and mediating locomotion behavior of the zebrafish embryo and larva. Development 123, 399–413.

Gu Z, Gu L, Eils R, Schlesner M & Brors B (2014). circlize Implements and enhances circular visualization in R. Bioinformatics 30, 2811–2812.

Guiraud S, Aartsma-Rus A, Vieira NM, Davies KE, van Ommen GJB & Kunkel LM (2015). The pathogenesis and therapy of muscular dystrophies. Annu Rev Genomics Hum Genet 16, 281–308.

Gupta VA, Ravenscroft G, Shaheen R, Todd EJ, Swanson LC, Shiina M, Ogata K, Hsu C, Clarke NF, Darras BT, Farrar MA, Hashem A, Manton ND, Muntoni F, North KN, Sandaradura SA, Nishino I, Hayashi YK, Sewry CA, Thompson EM, Yau KS, Brownstein CA, Yu TW, Allcock RJN, Davis MR, Wallgren-Pettersson C, Matsumoto N, Alkuraya FS, Laing NG & Beggs AH (2013). Identification of KLHL41 mutations implicates BTB-Kelch-mediated ubiquitination as an alternate pathway to myofibrillar disruption in nemaline myopathy. Am. J. Hum. Genet. 93, 1108–1117.

Guyon JR, Goswami J, Jun SJ, Thorne M, Howell M, Pusack T, Kawahara G, Steffen LS, Galdzicki M & Kunkel LM (2009). Genetic isolation and characterization of a splicing mutant of zebrafish dystrophin. Hum. Mol. Genet. 18, 202–211.

Hedges LV (1981). Distribution Theory for Glass’s Estimator of Effect Size and Related Estimators. Journal of Educational Statistics 6, 107–128.

Karatzoglou A, Smola A, Hornik K & Zeileis A (2004). kernlab – An S4 package for kernel methods in R. J Stat Softw 11, 1–20.

Kassambara A & Mundt F (2020). factoextra: Extract and visualize the results of multivariate data analyses https://CRAN.R-project.org/package=factoextra.

Kawahara G, Karpf JA, Myers JA, Alexander MS, Guyon JR & Kunkel LM (2011). Drug screening in a zebrafish model of Duchenne muscular dystrophy. Proc. Natl. Acad. Sci. U.S.A. 108, 5331–5336.

Khan MRA & Brandenburger T (2024). ROCit: performance assessment of binary classifier with visualization https://cran.r-project.org/package=ROCit.

Kirby KN & Gerlanc D (2013). BootES: an R package for bootstrap confidence intervals on effect sizes. Behav Res Methods 45, 905–927.

Koenig M, Hoffman EP, Bertelson CJ, Monaco AP, Feener C & Kunkel LM (1987). Complete cloning of the Duchenne muscular dystrophy (DMD) cDNA and preliminary genomic organization of the DMD gene in normal and affected individuals. Cell 50, 509–517.

Lambert MR & Gussoni E (2023). Tropomyosin 3 (TPM3) function in skeletal muscle and in myopathy. Skelet Muscle 13, 18.

Lambert MR, Spinazzola JM, Widrick JJ, Pakula A, Conner JR, Chin JE, Owens JM & Kunkel LM (2021). PDE10A inhibition reduces the manifestation of pathology in DMD zebrafish and represses the genetic modifier PITPNA. Mol Ther 29, 1086–1101.

Lê S, Josse J & Husson F (2008). FactoMineR: An R Package for Multivariate Analysis. Journal of Statistical Software 25, 1–18.

Li M, Andersson-Lendahl M, Sejersen T & Arner A (2014). Muscle dysfunction and structural defects of dystrophin-null sapje mutant zebrafish larvae are rescued by ataluren treatment. FASEB J. 28, 1593–1599.

Makowski D, Lüdecke D, Patil I, Thériault R, Ben-Shachar MS & Wiernik BM (2023). Automated results reporting as a practical tool to improve reproducibility and methodological best practices adoption. CRAN.

Marques JC, Lackner S, Félix R & Orger MB (2018). Structure of the zebrafish locomotor repertoire revealed with unsupervised behavioral clustering. Curr. Biol. 28, 181–195.e5.

Mathis A, Mamidanna P, Cury KM, Abe T, Murthy VN, Mathis MW & Bethge M (2018). DeepLabCut: markerless pose estimation of user-defined body parts with deep learning. Nat Neurosci 21, 1281–1289.

Meyer D, Dimitriadou E, Hornik K, Weingessel A, Leisch F, Chang CC & Lin CC (2023). e1071 https://CRAN.R-project.org/package=e1071.

Müller UK, van den Boogaart JGM & van Leeuwen JL (2008). Flow patterns of larval fish: undulatory swimming in the intermediate flow regime. J. Exp. Biol. 211, 196–205.

Müller UK & van Leeuwen JL (2004). Swimming of larval zebrafish: ontogeny of body waves and implications for locomotory development. J. Exp. Biol. 207, 853–868.

Nath T, Mathis A, Chen AC, Patel A, Bethge M & Mathis MW (2019). Using DeepLab-Cut for 3D markerless pose estimation across species and behaviors. Nat Protoc 14, 2152–2176.

Pedersen TL (2024). patchwork: The composer of plots https://CRAN.R-project.org/package=patchwork.

Pedersen TL & Robinson D (2024). gganimate: a grammar of animated graphics https://gganimate.com.

R Core Team (2024). R: a language and environment for statistical computing https://www.R-project.org/.

Robertson DGE, Caldwell GE, Hamill J, Kamen G & Whittlesey SN (2014). Research Methods in Biomechanics Human Kinetics, Champaign, IL, 2nd edition edition.

Robinson PN (2012). Deep phenotyping for precision medicine. Hum Mutat 33, 777–780.

Sheppard K, Gardin J, Sabnis GS, Peer A, Darrell M, Deats S, Geuther B, Lutz CM & Kumar V (2022). Stride-level analysis of mouse open field behavior using deep-learning-based pose estimation. Cell Rep 38, 110231.

Sing T, Sander O, Beerenwinkel N & Lengauer T (2005). ROCR: visualizing classifier performance in R. Bioinformatics 21, 3940–3941.

Smith LL, Beggs AH & Gupta VA (2013). Analysis of skeletal muscle defects in larval zebrafish by birefringence and touch-evoke escape response assays. J. Vis. Exp. 82, e50925.

Spinazzola JM, Lambert MR, Gibbs DE, Conner JR, Krikorian GL, Pareek P, Rago C & Kunkel LM (2020). Effect of serotonin modulation on dystrophin-deficient zebrafish. Biol Open 9, bio053363.

Stewart WJ & McHenry MJ (2010). Sensing the strike of a predator fish depends on the specific gravity of a prey fish. J Exp Biol 213, 3769–3777.

Tabor KM, Bergeron SA, Horstick EJ, Jordan DC, Aho V, Porkka-Heiskanen T, Haspel G & Burgess HA (2014). Direct activation of the Mauthner cell by electric field pulses drives ultrarapid escape responses. J. Neurophysiol. 112, 834–844.

Telfer WR, Nelson DD, Waugh T, Brooks SV & Dowling JJ (2012). Neb: a zebrafish model of nemaline myopathy due to nebulin mutation. Dis Model Mech 5, 389–396.

Ushey K (2018). RcppRoll: Efficient rolling / windowed operations https://CRAN.R-project.org/package=RcppRoll.

van den Brand T (2024). ggh4x: Hacks for ‘ggplot2’ https://CRAN.R-project.org/package=ggh4x.

van Leeuwen JL, van der Meulen T, Schipper H & Kranenbarg S (2008). A functional analysis of myotomal muscle-fibre reorientation in developing zebrafish Danio rerio. J. Exp. Biol. 211, 1289–1304.

van Raamsdonk W, van’t Veer L, Veeken K, Heyting C & Pool CW (1982). Differentiation of muscle fiber types in the teleost Brachydanio rerio, the zebrafish. Posthatching development. Anat. Embryol. 164, 51–62.

Wakeling JM & Johnston IA (1998). Muscle power output limits fast-start performance in fish. J. Exp. Biol. 201, 1505–1526.

Waugh TA, Horstick E, Hur J, Jackson SW, Davidson AE, Li X & Dowling JJ (2014). Fluoxetine prevents dystrophic changes in a zebrafish model of Duchenne muscular dystrophy. Hum. Mol. Genet. 23, 4651–4662.

Wickham H, Averick M, Bryan J, Chang W, McGowan LD, François R, Grolemund G, Hayes A, Henry L, Hester J, Kuhn M, Pedersen TL, Miller E, Bache SM, Müller K, Ooms J, Robinson D, Seidel DP, Spinu V, Takahashi K, Vaughan D, Wilke C, Woo K & Yutani H (2019). Welcome to the tidyverse. J Open Source Softw 4, 1686.

Widrick JJ, Alexander MS, Sanchez B, Gibbs DE, Kawahara G, Beggs AH & Kunkel LM (2016). Muscle dysfunction in a zebrafish model of Duchenne muscular dystrophy. Physiol. Genomics 48, 850–860.

Widrick JJ, Lambert MR, Kunkel LM & Beggs AH (2023). Optimizing assays of zebrafish larvae swimming performance for drug discovery. Expert Opin Drug Discov 18, 629–641.

Wilke CO (2024). cowplot: Streamlined plot theme and plot annotations for ‘ggplot2’ https://CRAN.R-project.org/package=cowplot.

Wilke CO & Wiernik BM (2022). ggtext: Improved text rendering support for ‘ggplot2’ https://CRAN.R-project.org/package=ggtext.

Xie Y (2024). knitr: a general-purpose package for dynamic report generation in R https://yihui.org/knitr/.

Zhu H (2024). kableExtra: Construct complex table with ‘kable’ and pipe syntax https://CRAN.R-project.org/package=kableExtra.

